# Inter and Intralaminar Excitation of Parvalbumin Interneurons in Mouse Barrel Cortex

**DOI:** 10.1101/2023.06.02.543448

**Authors:** Kate S. Scheuer, Anna M. Jansson, Xinyu Zhao, Meyer B. Jackson

## Abstract

Parvalbumin (PV) interneurons are inhibitory fast-spiking cells with essential roles in directing the flow of information through cortical circuits. These neurons set the balance between excitation and inhibition, control rhythmic activity, and have been linked to disorders including autism spectrum and schizophrenia. PV interneurons differ between cortical layers in their morphology, circuitry, and function, but how their electrophysiological properties vary has received little attention. Here we investigate responses of PV interneurons in different layers of primary somatosensory barrel cortex (BC) to different excitatory inputs. With the genetically-encoded hybrid voltage sensor, hVOS, we recorded voltage changes simultaneously in many L2/3 and L4 PV interneurons to stimulation in either L2/3 or L4. Decay-times were consistent across L2/3 and L4. Amplitude, half-width, and rise-time were greater for PV interneurons residing in L2/3 compared to L4. Stimulation in L2/3 elicited responses in both L2/3 and L4 with longer latency compared to stimulation in L4. These differences in latency between layers could influence their windows for temporal integration. Thus PV interneurons in different cortical layers of BC show differences in response properties with potential roles in cortical computations.

**Key points summary:**

- Excitatory synaptic responses were imaged in parvalbumin (PV) interneurons in slices of mouse barrel cortex using a targeted genetically-encoded voltage sensor. This approach revealed simultaneous voltage changes in approximately 20 neurons pre slice in response to stimulation.
- PV interneurons residing in layer 2/3 had larger amplitudes, longer half-widths, and longer rise-times than PV interneurons residing in layer 4.
- Responses of PV interneurons residing in either layer 2/3 or layer 4 had shorter latencies to stimulation of layer 4 compared to stimulation of layer 2/3.
- Excitatory synaptic transmission to PV interneurons varies with layer of residence and source of excitation.

## Introduction

Parvalbumin (PV) interneurons are inhibitory neurons defined by their expression of the calcium-binding protein PV (Tremblay *et al*., 2016). These fast-spiking cells are present in cortical layers 2-6 and play critical roles in controlling excitation/inhibition balance (Ferguson & Gao, 2018; Nahar *et al*., 2021), and in the generation of gamma oscillations, 30-80 Hz brain waves implicated in many functions including working memory, attention, and perceptual binding (Tallon-Baudry *et al*., 1998; Gonzalez-Burgos *et al*., 2015). PV interneurons and gamma oscillations have both been linked to a variety of psychiatric conditions including schizophrenia and autism spectrum disorder (Gonzalez-Burgos *et al*., 2015; Lauber *et al*., 2018; Kayarian *et al*., 2020).

Primary somatosensory barrel cortex (BC) is an attractive place to study PV interneurons because of its well-defined functions and architecture (Brecht, 2007; Feldmeyer, 2012; Feldmeyer *et al*., 2018; Staiger & Petersen, 2021). BC is defined by the presence of barrels, cytoarchitectural units in L4 which each correspond to a single vibrissa (Woolsey & Van der Loos, 1970). Distinct molecular, morphological, and electrophysiological cell types form complex circuits both within and between cortical layers in BC. Two major PV interneuron morphological subgroups, basket cells and chandelier cells, are distributed differently across cortical layers in BC. Chandelier cells form axoaxonic contacts and are not present in L4 (Li & Huntsman, 2014), while basket cells provide perisomatic inhibition and can be found in L2-6 (Naka & Adesnik, 2016; Frandolig *et al*., 2019; Staiger & Petersen, 2021). These two morphological subgroups can form distinct interlaminar circuits (Xu & Callaway, 2009). In addition to these morphological differences, PV interneurons in different layers may have different roles in functions such as intracortical and thalamic integration (Staiger & Petersen, 2021). Additionally, optogenetically generated gamma oscillations within a given cortical layer inhibit locally within that layer but facilitate in other layers. Furthermore, peak gamma oscillation power was higher for L6 compared to L2/3 (Adesnik, 2018). These results raise the important question of whether the roles of PV interneurons in different layers reflect differences in their responses to excitatory synaptic inputs. However a simultaneous assessment of voltage responses of PV neurons in different cortical layers has not been carried out.

Here we use the genetically-encoded hybrid voltage sensor (hVOS) to record excitatory post-synaptic potentials (EPSPs) optically from L2/3 and L4 PV interneurons in slices of mouse BC (Chanda *et al*., 2005; Wang *et al*., 2010; Bayguinov *et al*., 2017). We determined PV interneuron response amplitude, half-width, latency, rise-time, and decay-time elicited by stimulation in L2/3 and L4. Regardless of stimulation layer, L2/3 PV interneuron responses had higher amplitudes, longer rise-times, and broader half-widths than L4 PV interneurons. Additionally, responses to stimulation in L2/3 had longer latencies than responses to L4 stimulation, even after accounting for the effect of conduction distance. By contrast, responses in these layers had similar decay-times. Thus, hVOS imaging reveals variations in electrophysiological properties of PV interneurons between cortical layers. These differences have implications for how the cortex performs computations and integrates inputs between different layers.

## Methods

### Animals

PV-Cre driver mice (B6.129P2-Pvalb^tm1(cre)Arbr^/J, JAX strain 017320) were crossed with Ai35-hVOS1.5 Cre-reporter mice (C57BL/6-*Gt(ROSA)26Sor^tm1(CAG-hVOS1.5)Mbja^*/J, JAX strain 031102) to generate animals with hVOS probe targeted to PV interneurons (Bayguinov *et al*., 2017). Animal procedures were approved by the University of Wisconsin-Madison School of Medicine and Public Health Animal Care and Use Committee (IACUC protocol: M005952).

### hVOS probe

An hVOS probe was used to image voltage changes in PV interneurons. The probe used here is comprised of cerulean fluorescent protein (CeFP) tethered to the inner leaflet of the cell membrane with a truncated h-ras motif (Wang *et al*., 2010). Cells expressing the probe fluoresce, and this fluorescence is modulated by a Förster resonance energy transfer interaction with dipicrylamine (DPA), a small, hydrophobic anion which partitions into the cell membrane and moves when the membrane potential changes. Depolarization drives DPA towards the CeFP and fluorescence is quenched. Repolarization drives DPA back away from the CeFP so fluorescence increases. Fluorescence thus reports voltage changes of cells expressing the hVOS probe (Chanda *et al*., 2005; Wang *et al*., 2010). hVOS has sub-millisecond temporal resolution (Chanda *et al*., 2005; Bradley *et al*., 2009) and can be genetically targeted to specific cell types using a Cre-lox system (Bayguinov *et al*., 2017). Our crossing of hVOS Cre-reporter animals with PV Cre-driver animals produces mice previously shown to have 99.2% targeting specificity and express the hVOS probe in 83% of PV interneurons (Bayguinov *et al*., 2017).

### Slice preparation

Mice 7-8 weeks old were deeply anesthetized with isoflurane and sacrificed with cervical dislocation (institutional protocol noted above). Brains were rapidly dissected and placed into ice-cold cutting solution (in mM: 10 glucose, 125 NaCl, 4 KCl, 1.25 NaH_2_PO_4_, 26 NaHCO_3_, 6 MgSO_4_, 1 CaCl_2_) bubbled with a mixture of 95% O_2_ / 5% CO_2_. After approximately five minutes, brains were mounted and cut into 300 μm thick coronal slices with a with a Leica VT1200S vibratome. Slices were placed into a chamber filled with 95% O_2_ / 5% CO_2_-bubbled artificial cerebrospinal fluid (ACSF) with the same composition as cutting solution except with 1.3 mM MgSO_4_ and 2.5 mM CaCl_2_, and allowed to recover for at least 45 minutes.

### Electrophysiology and Imaging

Imaging experiments were performed in 95% O_2_ / 5% CO_2_-bubbled ACSF containing 4 μM DPA. Slices were placed into a custom recording chamber, and viewed with a BX51 Olympus microscope. Stimulus pulses 200 μA, 180 μsec were applied with a stimulus isolator (World Precision Instruments, Sarasota, Florida) through fire-polished, ACSF-filled KG-33 glass electrodes (King Precision Glass, Claremont, California) with tip diameters of about 6-8 μm. Stimulating electrodes were positioned in L2/3 or L4 of BC using a micromanipulator. Slices were illuminated with an LED with peak emission at 435 nm (Prizmatix, Holon, Israel) through a CFP filter cube. PV interneuron responses were acquired with a CCD-SMQ camera (RedShirt Imaging, Decatur, Georgia) at 2000 Hz with 80x80 spatial resolution. Bandpass filters of 5 and 10 nm centered at 435 nm were added to the excitation pathway when resting light intensities saturated the CCD-SMQ camera. Gradient contrast and higher resolution fluorescence images were captured by directing light to a Kiralux CMOS camera (Thorlabs, Newton, New Jersey). Data acquisition and analysis was performed with custom software (Chang, 2006).

### Identifying individual responsive PV interneurons

PV interneurons have extensive axonal and dendritic arbors which allow them to sample input from many cells, and to provide strong, widespread inhibition (Fukuda & Kosaka, 2003; Povysheva *et al*., 2008; Packer & Yuste, 2011; Hu *et al*., 2014). Cortical PV interneurons contact about 43-50% or more of pyramidal cells within about < 200 μm (Packer & Yuste, 2011; Inan *et al*., 2013), and fast-spiking interneuron to excitatory cell connectivity in BC can be as high as 67% for a sub-group of PV interneurons in L4 (Koelbl *et al*., 2015). Given the dense dendritic and axonal arbors of PV interneurons, a plasma membrane label such as the hVOS probe produces broad diffuse fluorescence throughout a slice, obscuring the fluorescence from PV interneuron cell bodies. This makes it difficult to identify individual PV interneuron cell bodies, despite clearly identifiable cortical layers in both gradient contrast (Fig. 1A) and fluorescence (Fig. 1B) images.

**Figure 1.**
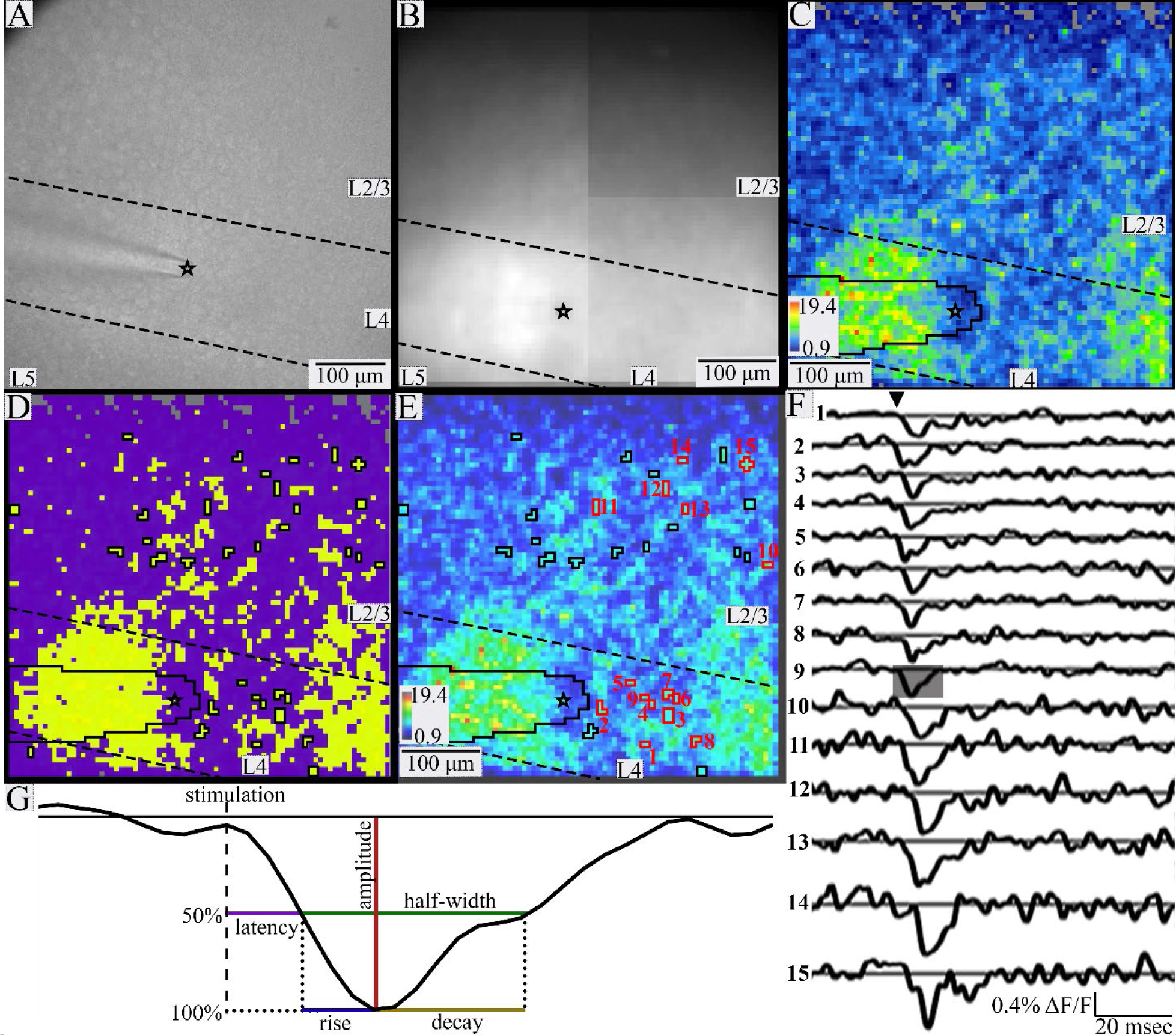
Identifying individual responsive PV interneurons. Gradient contrast image taken with the Kiralux camera (A) and fluorescence image taken with the CCD-SMQ camera (B) of a BC slice. Black stars indicate the tip of the stimulating electrode, and dashed lines indicate layer boundaries in A-E. C. SNR heatmap of slice in A and B. Gray pixels near the top have signals below the baseline noise and were excluded from analysis. The electrode is outlined in black on the lower left edge in C-E. D. K-means cluster map. K-means clustering of SNR was performed on pixels with SNR > baseline (colored in C). The data were best fitted with two clusters, with averages of 4.8 and 9.2. The yellow cluster with higher average SNR is likely to contain responsive PV interneurons, while the purple cluster with lower average SNR probably contains processes and unresponsive neurons. E. SNR heatmap overlaid with identified responsive PV interneurons outlined in black or red (color choice based on ease of view and not cell properties). F. Traces of fluorescence versus time for the PV interneurons outlined and numbered in E show clear depolarization in response to stimulation (triangle). G. Expanded portion of a trace (12 msec) shaded in F illustrating response parameters. Amplitude (red) is the maximum change in fluorescence; latency (purple) is the time from stimulation to half-maximal change in fluorescence; half-width (green) is the time between half-maximal change in fluorescence from depolarization to repolarization; rise-time (blue) is the time between half-maximal and maximal change in fluorescence; decay-time (gold) is the time from peak to half-maximal fluorescence.

To address this problem, we developed a semi-automated method of analyzing imaging data for objective, reproducible identification of responsive PV interneuron somata. This method relied on a hybrid approach using both geometric constraints and K-means clustering of signal-to-noise ratio (SNR) values. SNR was calculated as the peak stimulus-evoked fluorescence change divided by the baseline root-mean-square fluorescence in a 20-msec pre-stimulus time window. Pixels with a SNR below the baseline noise were discarded as clearly unresponsive (gray pixels near top, Fig. 1C). For the geometric constraints, we required regions of interest (ROIs) representing responsive PV interneurons to be spatially distinct, with no shared faces. Acceptable ROIs were groups of up to nine contiguous pixels three or fewer pixels across. With 6 μm pixel dimensions this constrains groups to the size of a murine PV interneuron soma, which is approximately 20 μm in diameter (Wang *et al*., 2002; Selby *et al*., 2007; Kooijmans *et al*., 2020). This criterion excluded some larger groups of pixels as potentially representing more than one cell even though they had very clear responses with a high SNR. For example, the red, orange, and yellow regions in the lower left and lower right corners of Fig. 1C and 1E had high SNR values but formed groups which were much larger than 3 pixels across and clearly contained several responsive cells. Pixels in these areas were excluded from analysis. Because a single pixel (6 μm) is too small to be a cell body and could contain several overlapping PV interneuron dendrites (<0.5-3 μm in rats, smaller in mice (Muller *et al*., 2005; Judak *et al*., 2022)) and/or axons (< 1 μm, (Stedehouder *et al*., 2019)), single isolated pixels were not considered to be responsive PV interneurons, again despite their high SNR. Responses from pixels obscured by the stimulating electrode (outlined in black, Fig. 1C-E) were also excluded. Finally, responses < 45 μm from the tip of the stimulating electrode were assumed to be the result of direct stimulation and excluded.

To refine and corroborate this procedure, ROIs corresponding to putative responsive PV interneurons satisfying the geometric constraints were subjected to one-dimensional K-means clustering of SNR values. One-dimensional K-means clustering was performed on pixels with SNR above baseline noise (all but the gray pixels in Fig. 1C). Clustering served two main purposes. First, it divided pixels into groups with similar SNR values. Pixel clusters with higher average SNR are more likely to contain cell bodies (yellow, Fig. 1D) while those with lower average SNR are more likely to contain small processes or lack responsive cells (purple, Fig. 1D). We therefore set a response SNR cutoff for K-means clusters to 5 and excluded pixels in clusters of pixels with average SNR < this cutoff (purple, Fig. 1D), as they likely contained processes or unresponsive cells. For acceptable clusters (average SNR > 5) we assumed that if pixels within a group satisfying the geometric requirements have SNR values in the same K-means cluster, they are likely to represent the same cell body. K-means clustering identified groups of geometrically associated pixels with similar SNR and assigned them to specific cells. This method basically compared each pixel to its neighbors and grouped them based on the likelihood they represent the same cell. In summary, this method was conservative, implementing multiple exclusion criteria to focus on small groups of pixels with similar SNR that represent distinct, spatially separated neurons.

Traces of fluorescence versus time from groups of pixels identified in this way were manually inspected to verify appropriate responses to stimulation (Fig. 1F). We implemented an additional cutoff based on amplitude, and discarded pixel groups with average ΔF/F < 0.1%. Responses with amplitudes more than 3 times the standard deviation above the mean value (> 1.165%) were also excluded. Such instances were very rare (4 PV interneurons total), and occurred in particularly dark areas or corners of the field of view where resting light was very low and dividing resulted in implausibly high values. Because our analysis compares PV interneuron properties based on cortical layer, occasional cells on a border between cortical layers were also excluded.

Despite the conservative nature of this analysis, our procedure identified an average of 21 responsive PV interneurons per slice. Although anatomical estimates of PV interneuron density vary widely, conservative estimates suggest that our 480x480 μm field of view may contain up to approximately 75 PV interneurons (Keller *et al*., 2018). Our numbers are generally well below this, supporting our procedure as a conservative method of identifying responsive somata. This method provided a reproducible, objective, and robust procedure to identify individual responsive PV interneurons. A test of validity is presented in Results (Fig. 2).

**Figure 2.**
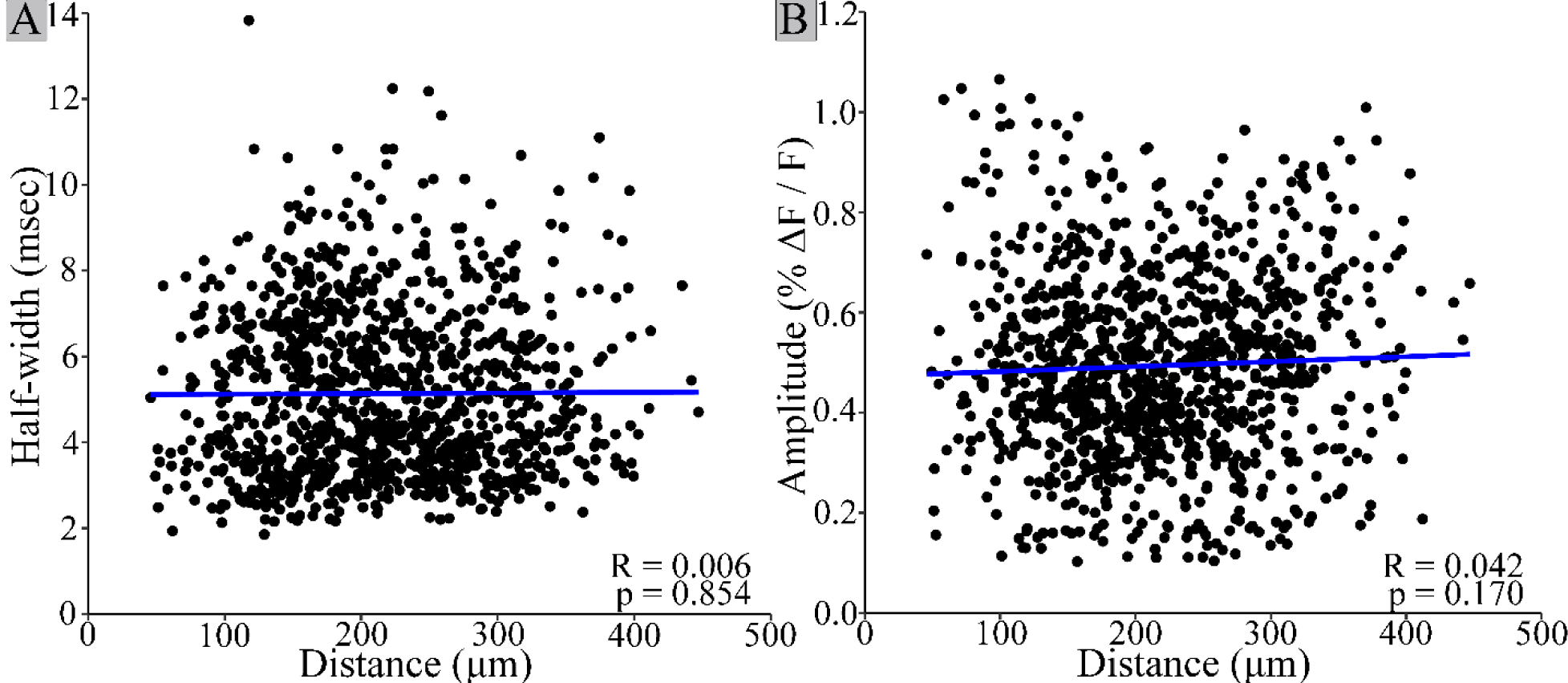
PV interneuron response half-width and amplitude do not vary with distance. Neither half-width (A, R = 0.006, p = 0.854) nor amplitude (B, Pearson’s product-moment correlation: R = 0.042, p = 0.170) are significantly correlated with distance from the stimulating electrode. This is consistent with single-cell responses, as half-width would be expected to increase, and amplitude would decrease with distance for a population response. Each point on the scatterplot corresponds to one PV interneuron. Linear regression best fit lines are shown in blue. N = 1086 cells from 52 slices.

Response parameters were extracted from traces of fluorescence versus time for pixel groups identified as corresponding to responsive PV interneurons (Fig. 1G). Amplitude is the maximum change in fluorescence. Latency is the time from stimulation to half-maximal change in fluorescence during depolarization. To account for the effect of distance on latency, we also divided the latency by distance to the stimulating electrode (distance-normalized latency). Half-width is the time between half-maximal change in fluorescence during depolarization and repolarization. Rise-time is the time from half-maximal to maximal fluorescence during depolarization. Decay-time is the time from maximum to half-maximal fluorescence during repolarization. All responses were examined visually and 4 were excluded because noise resulted in anomalous start times with obvious errors in parameters.

### Data processing and statistical tests

Fluorescence traces were processed with a nine-point binomial temporal filter and a spatial filter with σ = 1. A baseline determined from a polynomial fit was subtracted. Peak fluorescence change was divided by resting light intensity to give ΔF/F. Our method of responsive cell identification yielded 1086 PV interneurons from 52 slices from 7 animals (3 female, 4 male). Relationships between distance and half-width or amplitude were evaluated with Pearson’s product-moment correlation tests using individual PV interneurons as the unit of analysis. For remaining statistical tests, within a given slice layers with fewer than 8 responsive PV interneurons were excluded, values were averaged for all PV interneurons within each layer, and this average was used as the unit of analysis.

Normality was evaluated using Shapiro-Wilks tests. All parameters were normally distributed (amplitude: W = 0.982, p = 0.591; half-width: W = 0.986, p = 0.757; rise-time: W = 0.975, p = 0.300) or log-normally distributed (distance-normalized latency: W = 0.971, p = 0.210; decay-time: W = 0.986, p = 0.750). Variance between analysis groups was evaluated with Levene’s tests. Variance did not differ significantly for any parameter based on sex (amplitude: F(1,53) = 0.903, p = 0.346; half-width: F(1,53) = 0.078, p = 0.782; distance-normalized latency: F(1,53) = 3.808, p = 0.056; rise-time: F(1,53) = 0.353, p = 0.555; decay-time: F(1,53) = 0.252, p = 0.618). The effect of sex was evaluated with t-tests and showed no significant impact (amplitude: t(52.857) = 0.488, p = 0.628; half-width: t(50.648) = −1.018, p = 0.314; distance-normalized latency: t(52.438) = −1.252, p = 0.216; rise-time: t(48.567) = −0.652, p = 0.517; decay-time: t(52.005) = −0.893, p = 0.376).

Variance did not differ significantly based on PV interneuron layer or stimulation layer for any parameter (amplitude (F(3,51) = 0.465, p = 0.708); half-width (F(3,51) = 0.593, p = 0.623); distance-normalized latency (F(3,51) = 1.052, p = 0.378); rise-time (F(3,51) = 0.596 p = 0.321); decay-time (F(3,51) = 0.223, p = 0.880)). The effects of stimulation layer and/or PV interneuron residence layer on each parameter were therefore evaluated with ANOVA and post-hoc Tukey’s honestly significant differences tests.

### Code availability

R code, Python code, and custom software available on request.

## Results

### Validation of single cell identification

We tested our procedure of cell identification by examining variations with distance. If pixel groups actually represent multiple neighboring neurons rather than a single neuron, we would expect response half-width to broaden and response amplitude to decrease with distance from the stimulating electrode, as the activation of more distant groups should be less synchronous compared to closer groups. In plots versus distance neither parameter was significantly correlated with distance (Fig. 2, half-width: R = 0.006, p = 0.854; amplitude: R = 0.042, p = 0.170), indicating that pixel clusters do not contain more than one PV interneuron.

### PV interneuron responses vary between cortical layers

Stimulation in L2/3 (Fig. 3A-B, left) or L4 (Fig. 3A-B, right) elicited responses across L2/3 through L5 as shown in SNR heatmaps (Fig. 3C). Although PV interneurons in L5 responded to stimulation in either L2/3 or L4, less L5 was present in the field of view selected for study so we did not see enough L5 PV interneuron responses to include in the current analysis. L2/3 and L4 PV interneuron response parameters are presented in Table 1. Comparisons are presented in bar graphs below and will be discussed in detail.

**Figure 3.**
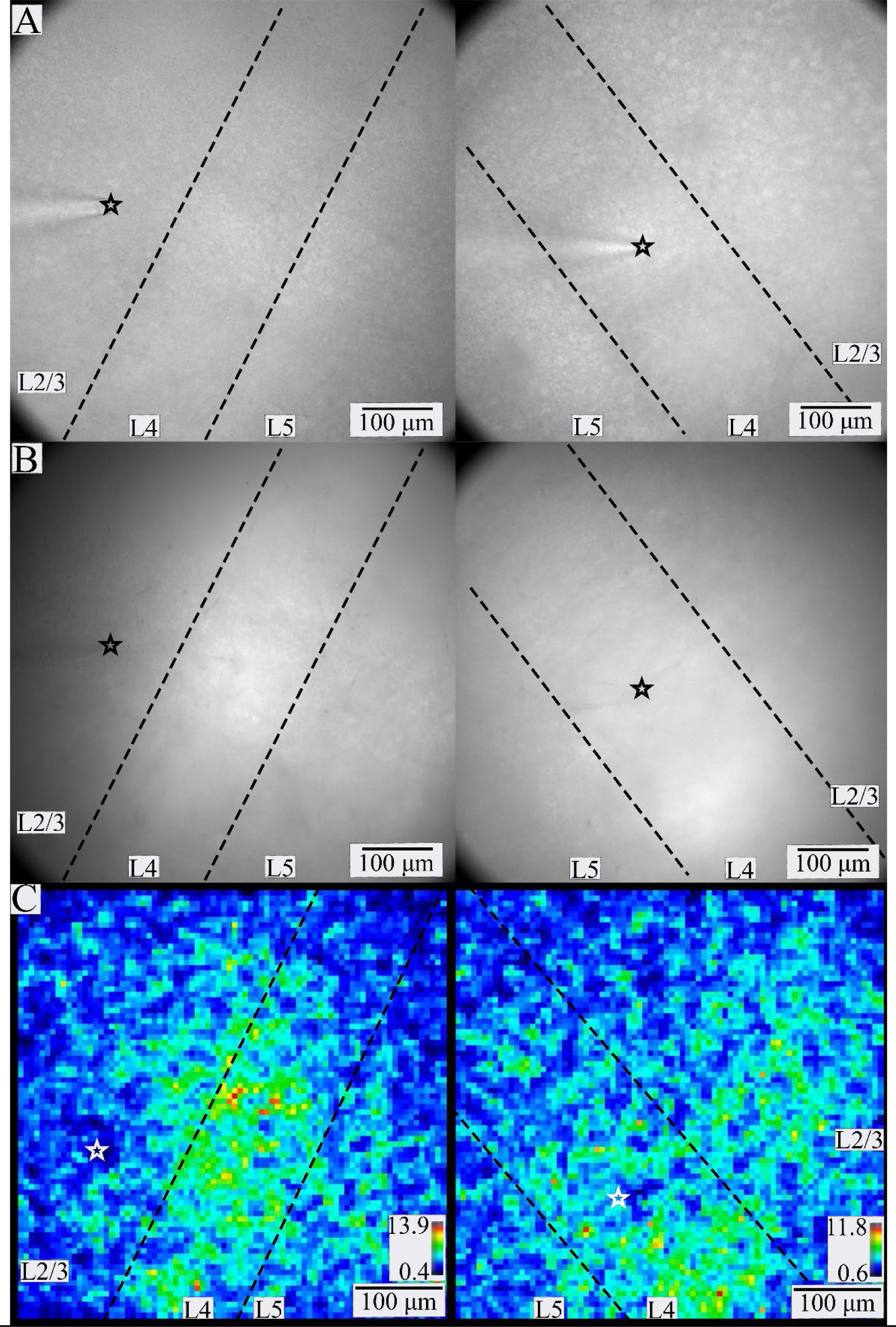
PV interneuron responses in BC. Gradient contrast (A) and fluorescence (B) images of two different slices of BC. L2/3 through L5 are visible within the fields of view. The tip of the stimulating electrode (black or white star) is visible in L2/3 (left) or L4 (right) in A-C. Dashed lines separate layers. C. SNR heatmaps for the slices shown in A and B. Warmer colors correspond to higher SNR regions more likely to contain responsive PV interneurons (color scales and ranges – lower right).

**Table 1.** PV interneuron response parameters based on residence layer and stimulation layer. Values are mean ± SE, N = number of neurons.

|  | L2/3 Stimulation |  | L4 Stimulation |  |
| --- | --- | --- | --- | --- |
|  | L2/3 PV<br>interneurons<br>(N = 322,<br>29 slices) | L4 PV<br>interneurons<br>(N = 259,<br>29 slices) | L2/3 PV<br>interneurons<br>(N = 331,<br>23 slices) | L4 PV<br>interneurons<br>(N = 160,<br>23 slices) |
| Amplitude (% $\Delta F/F$) | $0.567 \pm 0.005$ | $0.363 \pm 0.004$ | $0.556 \pm 0.005$ | $0.422 \pm 0.005$ |
| Half-width (msec) | $5.29 \pm 0.067$ | $4.82 \pm 0.049$ | $5.45 \pm 0.059$ | $4.71 \pm 0.050$ |
| Raw Latency (msec) | $2.79 \pm 0.059$ | $3.89 \pm 0.053$ | $3.28 \pm 0.050$ | $2.38 \pm 0.046$ |
| Distance-normalized<br>latency (msec/$\mu\text{m}$) | $0.015 \pm 0.0002$ | $0.017 \pm 0.0002$ | $0.013 \pm 0.0002$ | $0.014 \pm 0.0002$ |
| Rise-Time (msec) | $2.32 \pm 0.047$ | $2.17 \pm 0.039$ | $2.48 \pm 0.044$ | $2.02 \pm 0.039$ |
| Decay-Time (msec) | $2.97 \pm 0.051$ | $2.64 \pm 0.036$ | $2.97 \pm 0.043$ | $2.69 \pm 0.037$ |

PV interneuron residence layer significantly impacted amplitude (Fig. 4A, F(1,51) = 32.438, p < 0.001), rise-time (Fig. 4B, F(1,51) = 4.753, p = 0.034), and half-width (Fig. 4C, F(1,51) = 4.710, p = 0.035), but not decay-time (Fig. 4D, F(1,51) = 1.422, p = 0.239). Regardless of stimulation layer, PV interneuron response amplitudes (0.579 ± 0.014, mean ± SE) were 46% larger in L2/3 than in L4 (0.396 ± 0.018, mean ± SE, p < 0.001). Rise-times for PV interneurons residing in L2/3 (2.39 ± 0.070 msec) were also longer than those in L4 (2.13 ± 0.047 msec, p = 0.034). L2/3 PV interneuron response half-widths (5.26 ± 0.108 msec) were broader than L4 response half-widths (4.82 ± 0.091 msec, p = 0.035).

**Figure 4.**
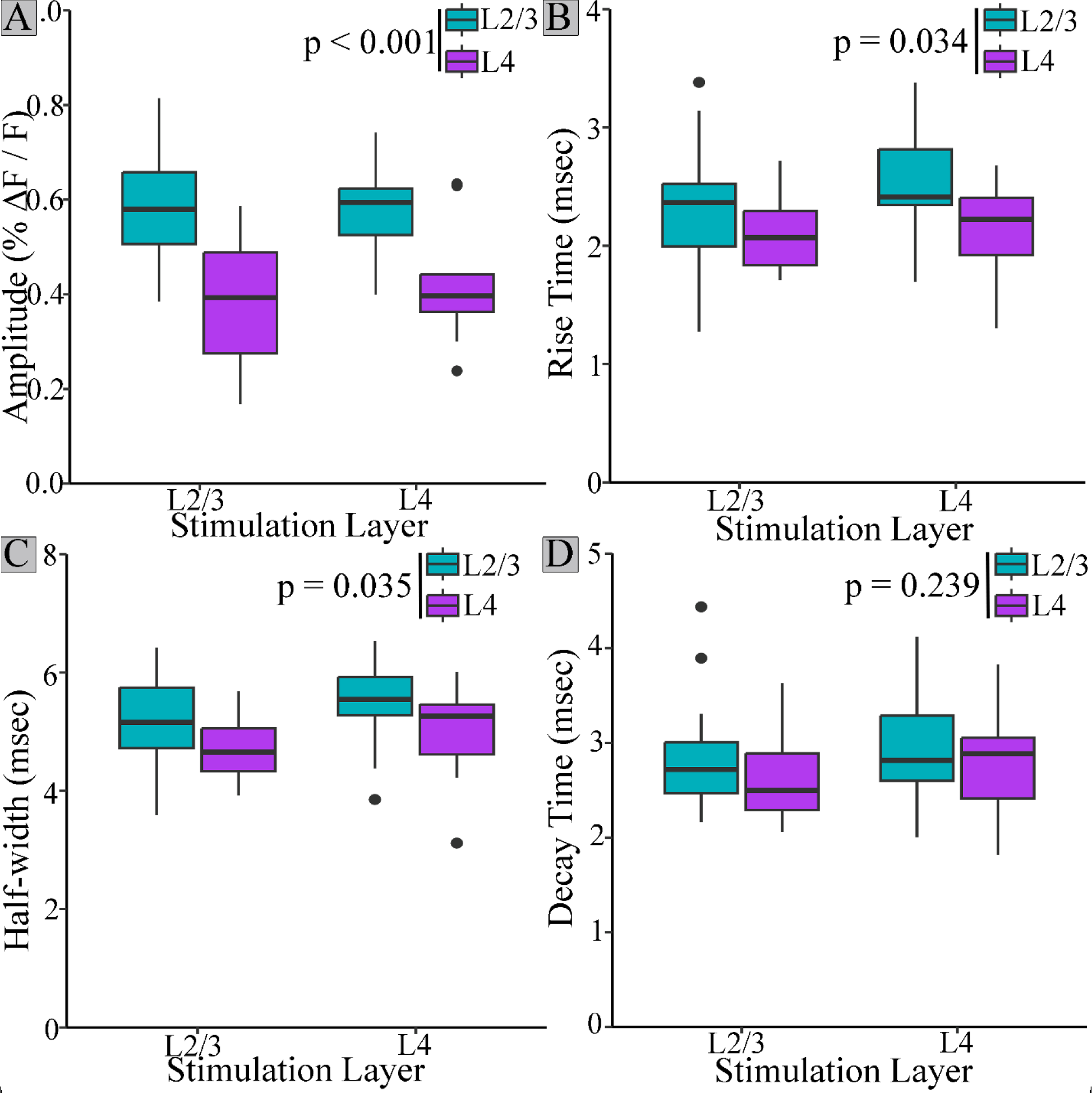
Amplitude, rise-time, and half-width of PV interneurons residing in different layers. L2/3 PV interneuron responses (blue) had higher amplitudes (A), longer rise-times (B), and broader half-widths (C) than responses from PV interneurons in L4 (purple). Decay-time (D) did not differ based on PV interneuron residence layer. Stimulation layer did not significantly impact amplitude, half-width, rise-time, or decay-time.

Latency also differed based on cortical layer (Fig. 5). Unlike amplitude, rise-time, and half-width, however, latency also depended on stimulation layer. To analyze variations in latency we must take into account the influence of distance and conduction time (Scheuer *et al*., 2023). Distances from the electrode tip to PV interneurons responding to L2/3 (180 ± 5.3 μm) and L4 (178 ± 4.3 μm) intralaminar stimulation were similar, as were distances for interlaminar responses to L2/3 (237 ± 3.4 μm) and L4 (260 ± 4.1 μm) stimulation. However, average interlaminar distance (248 ± 4.0 μm) was 39% longer than intralaminar distance (179 ± 4.9 μm). As expected, distance significantly impacted raw latency (F(1,53) = 49.62, p < 0.001). To account for this effect, we compared response latencies divided by distance from the tip of the stimulating electrode (distance-normalized latency). Stimulation layer significantly affected distance-normalized latency (F(1,51) = 16.478, p < 0.001). Regardless of PV interneuron layer, distance-normalized latencies of responses to stimulation in L2/3 (0.0162 ± 0.0005 msec/μm) were significantly longer than those to stimulation in L4 (0.0128 ± 0.0004 msec/μm, p < 0.001). PV interneurons residing in L2/3 responded to interlaminar L4 stimulation (0.013 ± 0.0003 msec/μm) more quickly than intralaminar L2/3 stimulation (0.016 ± 0.0004 msec/μm, t(26.992) = 3.243, p = 0.003). However, L4 PV interneurons responded to interlaminar L2/3 stimulation (0.017 ± 0.0005 msec/μm) more slowly than to intralaminar L4 stimulation (0.013 ± 0.0005 msec/μm). Thus, stimulation layer impacts latency in a manner which cannot be attributed to differences in distance.

**Figure 5.**
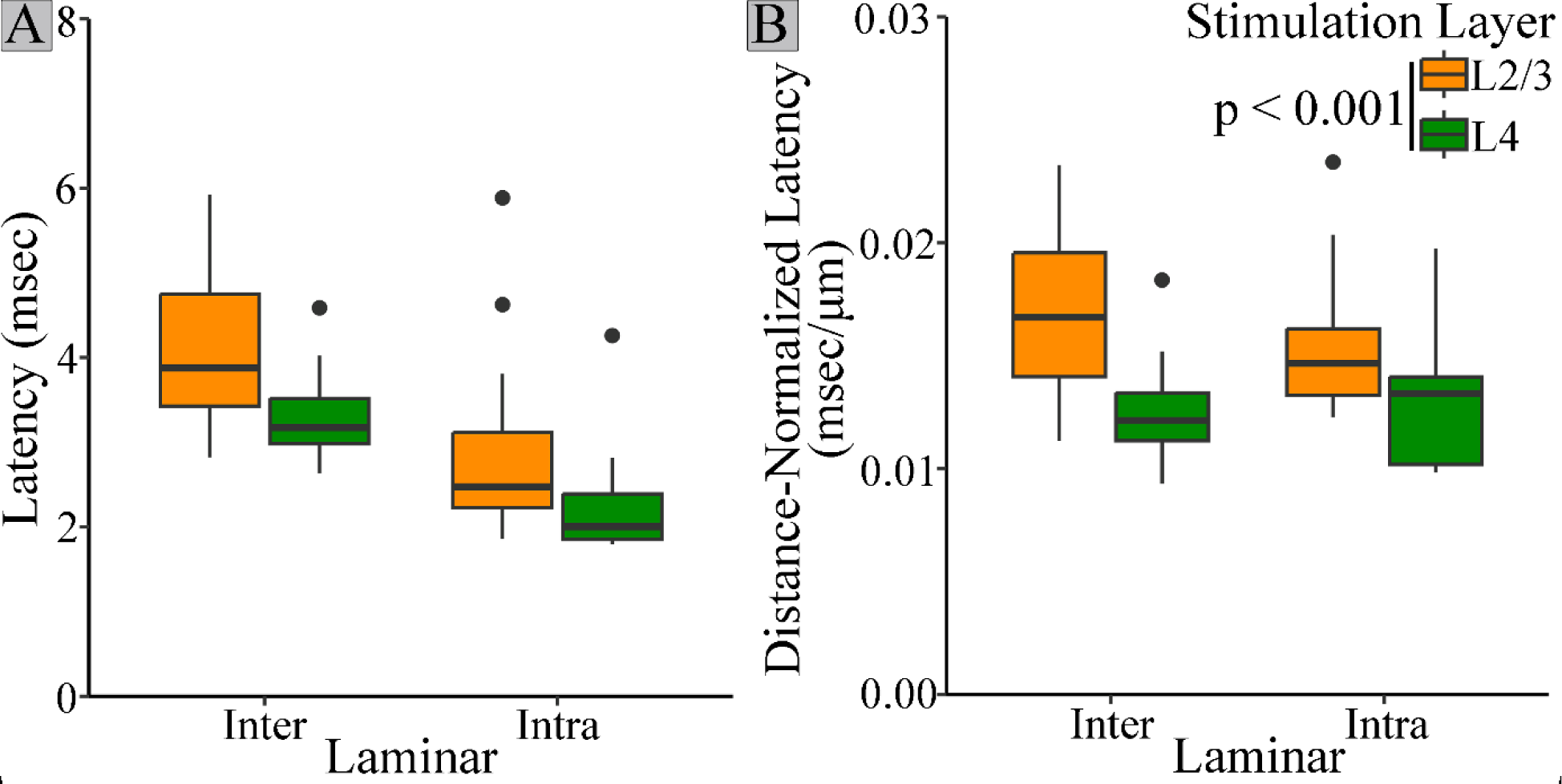
PV interneuron response latency depends on stimulation layer. A. Raw latencies for PV interneuron responses to stimulation in L2/3 (orange) or L4 (green). B. Regardless of residence layer, responses elicited by L2/3 stimulation (orange) have longer distance-normalized latencies than those elicited by L4 stimulation (green, t = (50) = 4.244, p < 0.001).

## Discussion

Here we used the genetically-encoded hybrid voltage sensor hVOS to investigate PV interneurons in BC. We observed responses of L2/3 and L4 PV interneuron to stimulation of both these layers. We developed a semi-automatic method of identifying individual responsive PV interneurons that combines geometric considerations with statistical K-means clustering of SNR. This method reliably located responsive cells within the high background fluorescence produced by the extensive arborization of PV interneurons. Using this method, we were able to identify cell bodies of ∼20 responsive PV interneurons per slice, and determine response amplitude, half-width, latency, rise-time, and decay-time of their responses to synaptic excitation. To test our method, we plotted half-width and amplitude versus distance from the electrode tip (Fig 2). The absence of correlations indicates that our pixel clusters contain single cells rather than multiple cells. The parameters reported in this study represent basic elements of cortical circuitry that may be useful in the development of accurate computational models designed to recapitulate the roles of fast-spiking interneurons in BC microcircuit computations (Avermann *et al*., 2012), and in the generation of synchronous activity (Di Garbo *et al*., 2004; Pervouchine *et al*., 2006).

Previous hVOS studies of PV interneuron activity in somatosensory cortex determined that spike-like responses similar to action potentials had a peak ΔF/F of 2.4% (Bayguinov *et al*., 2017), while unitary synaptic responses elicited by an action potential in a single excitatory neuron ranged from 0.2 – 0.4% (Canales *et al*., 2022). Assuming ∼100 mV action potentials, the mean amplitude of 0.494% reported here can be estimated as approximately 21 mV. The responses reported here are 64% larger than unitary excitatory responses and about one fifth the amplitude of spike-like responses. They are clearly too small to be action potentials, and are likely to represent EPSPs elicited by an average of about two excitatory neurons. Consistent with our assessment that these responses are synaptic potentials, our half-widths of ∼5.14 msec are 3.6 times broader than half-widths of PV interneuron spikes recorded with hVOS (Bayguinov *et al*., 2017). Our half-widths fall in the range of other studies of subthreshold, synaptic responses of PV interneuron in murine cortex of 4.6 to 22.3 msec (Thomson, 1997; Angulo *et al*., 1999; Thomson *et al*., 2002; Beierlein *et al*., 2003; Holmgren *et al*., 2003; Ali & Nelson, 2006; West *et al*., 2006; Ali *et al*., 2007; Avermann *et al*., 2012; Zhou & Roper, 2014). Our 2.8 msec half-decay-time corresponds to an exponential decay-time of 4.1 msec, which is within the range (3.5 to 12 msec) of previously reported values for EPSPs in murine cortical PV interneurons (Povysheva *et al*., 2006; Otsuka & Kawaguchi, 2009; Zaitsev & Lewis, 2013; Athilingam *et al*., 2017). Thus, parameter values reported here fall within the range of previous reports, and reveal how PV interneuron properties vary depending on location and source of excitation.

Response amplitude, distance-normalized latency, rise-time, and half-width differed based on cortical layer, and Fig. 6 illustrates the key differences. Regardless of stimulation layer, response amplitude was greater for PV interneurons in L2/3 compared to L4. EPSP amplitude depends on a wide variety of factors including number of inputs, dendritic location, and ion channel and receptor makeup (Ali & Nelson, 2006; Bonsi *et al*., 2007; Otsuka & Kawaguchi, 2009; Zaitsev *et al*., 2012; Pala & Petersen, 2018; Guo *et al*., 2020; Das *et al*., 2021). The increased EPSP amplitude in L2/3 compared to L4 PV interneurons may therefore reflect some of these factors. PV interneuron EPSP amplitudes are more than twice as large if the presynaptic excitatory cell and PV interneuron are reciprocally connected (Zaitsev & Lewis, 2013). Thus, the larger EPSP amplitudes of L2/3 PV interneurons could be related to higher reciprocal connectivity. Calcium-permeable AMPA receptors have also been shown to impact fast-spiking interneuron synaptic responses (Bonsi *et al*., 2007; Zaitsev *et al*., 2012), and differences in their distribution between layers could contribute to the present findings.

**Figure 6.**
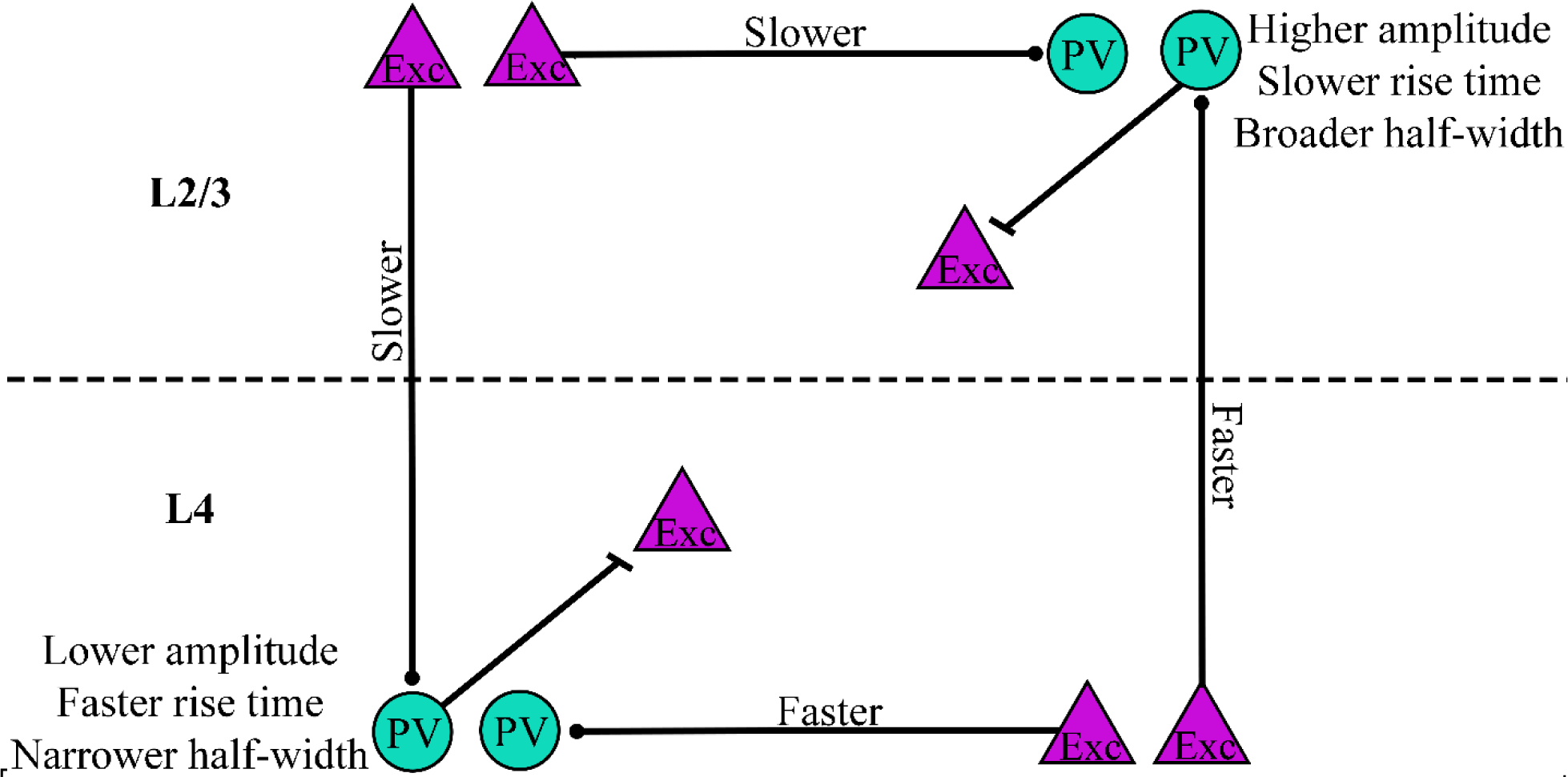
Summary of PV interneuron response differences. Amplitude, distance-normalized latency, rise-time, and half-width vary based on cortical layer. PV interneurons (teal circles) residing in L2/3 had higher amplitudes, slower rise-times, and broader half-widths compared to those in L4. Distance-normalized latencies of responses to stimulation of excitatory cells (purple triangles) in L2/3 were longer than those of responses to stimulation in L4.

PV interneuron residence layer also impacted rise-time and half-width. L2/3 PV interneuron responses had longer rise-times and broader half-widths than responses of PV interneurons in L4. Differences in rise-time could reflect dendritic location, presynaptic release kinetics, or AMPA receptor subunit composition. Compared to other types of interneurons and pyramidal cells, PV interneurons in CA1 express higher levels of AMPA receptor subunit GluA1, higher levels of auxiliary proteins regulating AMPA receptors, and especially high levels of GluA4 (Yamasaki *et al*., 2016). Knockout of GluA4, but not GluA3, decreases rise-time (Yang *et al*., 2011). AMPA receptors on PV interneurons can have multiple subunit combinations (Kondo *et al*., 1997). Therefore, differences in rise-time based on PV interneuron residence layer might reflect layer-specific variation in AMPA receptor subunit composition. Longer rise-times for PV interneurons in L2/3 compared to L4 likely contribute to the broader half-widths of L2/3 PV interneurons.

In addition to differences in amplitude, rise-time, and half-width, we also observed differences in latency between cortical layers. However, while the former properties depended on the layer in which PV interneurons resided, latency differed depending on the location of the inputs. Accounting for the effect of distance, both L2/3 and L4 PV interneuron responses to L2/3 stimulation had significantly longer latencies compared to responses to L4 stimulation. Because stimulation layer impacted latencies regardless of the layer in which the PV interneurons resided, this difference may reflect a property of the excitatory input rather than postsynaptic properties of PV interneurons. The inputs could have a faster conduction time along their axons or more direct axonal paths to their targets. It is also possible that the latency differences reflect differences in the kinetics of neurotransmitter release between axons originating in L2/3 and L4.

The shorter latencies of L2/3 PV interneuron responses to interlaminar L4 excitation compared to the reverse pathway of L4 PV interneuron responses to interlaminar L2/3 excitation will enable feedforward excitation along the canonical route from L4 to L2/3 PV interneurons to occur more quickly than L2/3 to L4 feedback. Inhibition plays a key role in coincidence detection by controlling the temporal integration window (Pouille & Scanziani, 2001), and because PV interneurons fire rapidly, they are particularly well-suited to tightly constraining integration within their synaptic targets. Compared to L4, which processes more basic sensory information such as touch and whisking, L2/3 functions are related to more complex somatosensory processing such as object localization (O’Connor *et al*., 2010), stimulus-specific adaptation (Yarden *et al*., 2022), texture discrimination (Allitt *et al*., 2017), and social touch (Lenschow & Brecht, 2015). The particularly rapid L4 to L2/3 excitation reported here could narrow the integration window set by L2/3 PV interneurons on their targets, and therefore may impact these higher-level sensory processes.

This work demonstrates the utility of hVOS voltage imaging as a technique to examine cortical circuitry of many cells of a specific type simultaneously across multiple cortical layers. This approach can be used to measure response parameters such as amplitude, half-width, latency, rise-time, and decay-time, which are important for computations. It also provides an opportunity to compare these response parameters across cortical layers. Here we observed layer-based differences in amplitude, rise-time, and latency which hold implications for how BC integrates interlaminar and intralaminar inputs. Future work building on this approach has the potential to address the circuit functions of PV interneurons as well as other specific cell types throughout the brain.

## Author contributions

Data collection (KSS), data analysis (KSS, AMJ), manuscript preparation (KSS, MBJ, XZ), project conception (MBJ, KSS, XZ)

## Conflict of interest statement

The authors declare no competing financial or non-financial interests.

## Acknowledgements

National Institutes of Health Grants NS127219 and NS093866 to MBJ and NS105200 XZ. Thanks to Dr. Shane McMahon for methodological contributions.

## Notes

### Competing Interest Statement

The authors have declared no competing interest.

### Summary of Updates

none

